# Assessment of Inactivation Procedures for SARS-CoV-2

**DOI:** 10.1101/2020.05.28.120444

**Authors:** Heidi Auerswald, Sokhoun Yann, Sokha Dul, Saraden In, Philippe Dussart, Nicholas J. Martin, Erik A. Karlsson, Jose A. Garcia-Rivera

**Author notes:** For correspondence on COVID-19 and other emerging infectious disease diagnostic testing and response in Southeast Asia, please contact Dr. Erik Karlsson. For correspondence on diagnostic testing for infectious disease on naval vessels and other austere sites, please contact Dr. Jose A. Garcia-Rivera.

## Abstract

Severe Acute Respiratory Syndrome Coronavirus 2 (SARS-CoV-2), the causative agent of Coronavirus disease 2019 (COVID-19), presents a challenge to laboratorians and healthcare workers around the world. Handling of biological samples from individuals infected with the SARS-CoV-2 virus requires strict biosafety and biosecurity measures. Within the laboratory, non-propagative work with samples containing the virus requires, at minimum, Biosafety Level-2 (BSL-2) techniques and facilities. Therefore, handling of SARS-CoV-2 samples remains a major concern in areas and conditions where biosafety and biosecurity for specimen handling is difficult to maintain, such as in rural laboratories or austere field testing sites. Inactivation through physical or chemical means can reduce the risk of handling live virus and increase testing ability worldwide. Herein we assess several chemical and physical inactivation techniques employed against SARS-CoV-2 isolates from Cambodian COVID-19 patients. This data demonstrates that all chemical (AVL, inactivating sample buffer and formaldehyde) and heat treatment (56°C and 98°C) methods tested completely inactivated viral loads of up to 5 log_10_.

## Introduction

Severe Acute Respiratory Syndrome Coronavirus 2 (SARS-CoV-2), the causative agent of Coronavirus disease 2019 (COVID-19) has rapidly spread across the world. On January 30^th^, 2020, the World Health Organization (WHO) declared the outbreak a Public Health Emergency of International Concern and upgraded it to a pandemic on March 11, 2020 [1]. As of May 28^th^, 2020, there have been over 5.6 million laboratory-confirmed cases and greater than 350,000 deaths reported globally [2]. Extensive testing is necessary to ensure accurate diagnosis, contact tracing, and mitigate SARS-CoV-2 spread through isolation and quarantine procedures. Additionally, extensive testing facilitates the global public health response against COVID-19, providing critical information regarding the effectiveness of mitigation efforts.

Given the lack of approved drugs and a vaccine, SARS-CoV-2 isolates should be handled according to strict biosafety and biosecurity measures [3, 4]. Extreme care in handling live samples prevents occupational exposure and requires extensive technical training and appropriate primary and secondary containment devices wearing recommended personal protective equipment. Therefore, with the need for extensive testing in areas and conditions where biosafety and biosecurity for specimen handling is difficult to maintain remains a major concern. Such scenarios include the need to sample outside of designated testing centers, conducting field investigations in difficult locales, non-secure sample transportation, and even testing in underequipped or under-maintained laboratories. Inactivation through physical or chemical means reduces the risk from handling live samples and increase testing ability worldwide.

Cambodia is a tropical, resource poor, least developed country (LDC) in Southeast Asia with a large socio-economic dependence on tourism [5]. Cambodia is also a major hotspot of endemic and emerging infectious disease [6]. One particular, but not unique, issue faced in LDCs is the expansion of testing capacity due to a scarcity of testing laboratories, especially in remote provincial health centers. Therefore, SARS-CoV-2 samples from these rural health centers requires safe but rapid transportation to designated testing sites. Aside from active training and great care when handling live specimens from suspected cases, transportation of potentially infectious material requires increased protective equipment and packaging, often in reduced supply or of poor quality, even in the best of scenarios. Therefore, simple and effective inactivation of suspected samples that can be conducted onsite can greatly decrease risk of exposure during transportation, handling, and testing, as well as reduce demand for protective equipment and supplies at a current global scarcity.

Herein, we evaluated the efficiency of various thermal and chemical inactivation methods on SARS-CoV-2 utilizing three separate SARS-CoV-2 isolates cultured from patient samples collected in Cambodia to determine their effect on viral infectivity and RNA integrity tested via real-time RT-PCR.

## Materials and Methods

### Cell lines

African green monkey kidney cells (Vero; ATCC# CCL-81) were used for the isolation and culture of SARS-CoV-2 isolates. Vero E6 cells were used for the titration of infectious virus via TCID_50_. Both cell lines were cultivated in Dulbecco’s modified Eagle medium (DMEM; Sigma-Aldrich, Steinheim, Germany) supplemented with 10% fetal bovine serum (FBS; Gibco, Gaithersburg, MD, USA) and 100 U/ml penicillin-streptomycin (Gibco) at 37°C and 5% CO_2_ atmosphere. Upon infection with SARS-CoV-2 the culture medium was replaced by infection medium containing DMEM, 5% FBS, antibiotics, 2.5 μg/mL Amphotericin B (Gibco) and 16 μg/mL TPCK-trypsin (Gibco).

### Virus culture and titration

Three SARS-CoV-2 isolates (designated hCoV-19/Cambodia/1775/2020, 1775; hCoV-19/Cambodia/2018/2020, 2018; and, hCoV-19/Cambodia/2310/2020, 2310) were obtained from patient’s swabs (combination of one nasopharyngeal and one oral swab in one tube of viral transport medium) and cultured in Vero cells. Virus-containing supernatants, as determined by the presence of cytopathic effect (CPE), were harvested six days after infection by centrifugation at 1,500 rpm for 10 min. The concentration of viable virus was measured by TCID_50_ assay on Vero E6 cells in 96-well plates (TPP, Trasadingen, Switzerland) [7]. Briefly, serial dilutions of viral culture supernatant were inoculated onto cells using infection medium. After 4 days of incubation, plates were inactivated with 4% formaldehyde for 20 minutes then stained with 1% crystal violet solution in phosphate buffered saline (PBS) for 20 min. Titer of viable virus was calculated applying the Spearman-Karber formula [8].

### Inactivation of SARS-CoV-2 isolates

Inactivation was performed in triplicate using 140 μL aliquots of SARS-CoV-2 isolates (1775, 2018, and 2310; passage 3 from Vero cells). Chemical inactivation included: (i) adding 560 μL viral lysis buffer (AVL buffer including carrier RNA; AVL buffer) from the QIAamp Viral RNA Mini Kit (Qiagen, Hilden, Germany) for 10 min at room temperature according the manufacturer’s recommendations; (ii) 200 μL inactivating sample buffer (GeneReach, Taichung City, Taiwan) containing 50% guanidinium thiocyanate (GITC) and 6% t-Octylphenoxypolyethoxyethanol (Triton X-100) for 15 min at room temperature; or, (iii) 140 μL 4% Formaldehyde in PBS (General Drugs House Co. Ltd., Bangkok, Thailand) for 15 min at room temperature. Thermal inactivation similarly performed on 140 μL aliquots of fresh virus culture: (iv) 56°C for 30 min; (v) 56°C for 60 min; and, (vi) 98°C for 2 min in a thermo-block (Eppendorf, Hamburg, Germany). Sterile DMEM treated in the similar methods served as negative controls, and untreated SARS-CoV-2 isolates as positive controls.

### Analysis for viable virus post inactivation

To determine if any viable virus remained post inactivation, 50% Polyethylene glycol 8000 (Sigma-Aldrich, St. Louis, USA) in PBS was added (1/5 of total sample volume) to an aliquot from each sample condition and incubated overnight at 4°C. Following incubation, virus was recovered by centrifugation at 1,500 rpm for 1h. Precipitates were washed twice with sterile PBS, re-constituted with infection medium, and used for infecting the TCID_50_ on Vero E6 cells and recovery cultures on Vero cells. Negative controls were treated the same way to examine cytotoxicity of possible remaining traces of inactivation solutions.

### SARS-CoV-2 real-time RT-PCR

Following inactivation, RNA from one aliquot per condition per virus isolate and negative control was immediately extracted with the QIAamp Viral RNA Mini Kit (Qiagen) and stored at −80°C until further processing. Real-time RT-PCR assays for SARS-CoV-2 RNA detection were performed in duplicate using the Charité Virologie algorithm (Berlin, Germany) to detect both E and RdRp genes [9]. In brief, real-time RT-PCR was performed using the SuperScript™ III One-Step RT-PCR System with Platinum™ Taq DNA Polymerase (Invitrogen) on the CFX96 Touch Real-Time PCR Detection System (BioRad, Hercules, CA, USA)ABI. Each 25 μl reaction mixture contained 5 μl of RNA, 3.1 μl of RNase-free water, 12.5 μl of 2X PCR buffer, 1 μl of SuperScriptTM III RT/Platinum Taq Mix, 0.5ul of each 10 μM forward and reverse primer, and 0.25 μl of probe (E_Sarbeco_P1 or RdRP_SARSr-P2) using the following thermal cycling conditions; 10 min at 55°C for reverse transcription, 3 min at 94°C for PCR initial activation, and 45 cycles of 15 s at 94°C and 30 s at 58°C.

### Statistical Analysis

Statistics were performed using GraphPad Prism for Windows, version 7.02 (GraphPad Software, Inc., La Jolla, CA,, USA). Analysis of variance was performed comparing mean Ct values for each inactivation method. Difference between standard (AVL) and each specific inactivation method was determined using Dunnett’s test for many-to-one comparison. A p-value of less than 0.05 was considered to indicate statistical significance. Agreement, including bias and 95% confidence interval, between Ct values following inactivation by AVL and other methods was assessed using a method described by Bland and Altman [10]. The mean Ct value of AVL and the other inactivation method assessed was plotted on the X-axis. The difference between the two values was plotted on the Y-axis. Cut-off values of 2 and − 2 are plotted.

## Results

### Inactivation efficiency

All chemical and thermal inactivation methods resulted in the reduction of viable SARS-CoV-2 to undetectable levels. Untreated virus isolates had a concentration of viable virus up to 6.67 × 10^5^ (isolate 2310) before treatment (Table 1). Therefore, the reduction of viable virus across inactivation levels was at least 5 log_10_. Precipitation of virus and complete removal of inactivation solution before infecting Vero E6 cells for TCID_50_ titration ensured that no CPE was induced by chemical products used in the inactivation procedure. Successful recovery of virus post-PEG precipitation was determined by RT-PCR on the same samples used for TCID_50_. All attempts to recover viable virus post inactivation on Vero cells were unsuccessful up to day 6 post-inoculation.

**Table 1:** SARS-CoV-2 isolates used for inactivation.

| **SARS-CoV-2 isolate** | **Before treatment**<br>**TCID<sub>50</sub>/mL** | **After PEG precipitation**<br>**(positive control)**<br>**TCID<sub>50</sub>/mL** | **Post inactivation**<br>**(all methods)**<br>**TCID<sub>50</sub>/mL** |
| --- | --- | --- | --- |
| 1775 | 2.11E+05 | 4.10E+04 | Not detected |
| 2018 | 1.19E+04 | 6.09E+03 | Not detected |
| 2310 | 6.67E+05 | 1.22E+05 | Not detected |

### Effect of inactivation procedure on RT-PCR

There were significant differences between the Ct values for the RdRp (ANOVA; p<0.0001) and E (ANOVA; p<0.0001) genes. Following many-to-one comparison of AVL to all other forms of inactivation used in this study (Figure 1), only formaldehyde inactivation was significantly different for the RdRp (Dunnet’s Test; p=0.0016) and E (Dunnet’s Test; p=0.0007) genes. In order to demonstrate the agreement in Ct values for the inactivation methods compared to the standard AVL, Bland-Altman plots are graphically presented in Figure 2 with cut-off values marked at two Ct differences (dashed lines). Samples inactivated by formaldehyde were the only ones where the absolute bias Ct value for all samples was greater than two compared to AVL for RdRp (−20.32 ± 1.75) and E (−19.80 ± 1.17) genes. All other inactivation methods resulted in absolute bias Ct values of less than one except for the RdRp gene following inactivation at 56°C for 30 min (−1.15 ± 1.08), though this was still within the limits of agreement.

**Figure 1:**
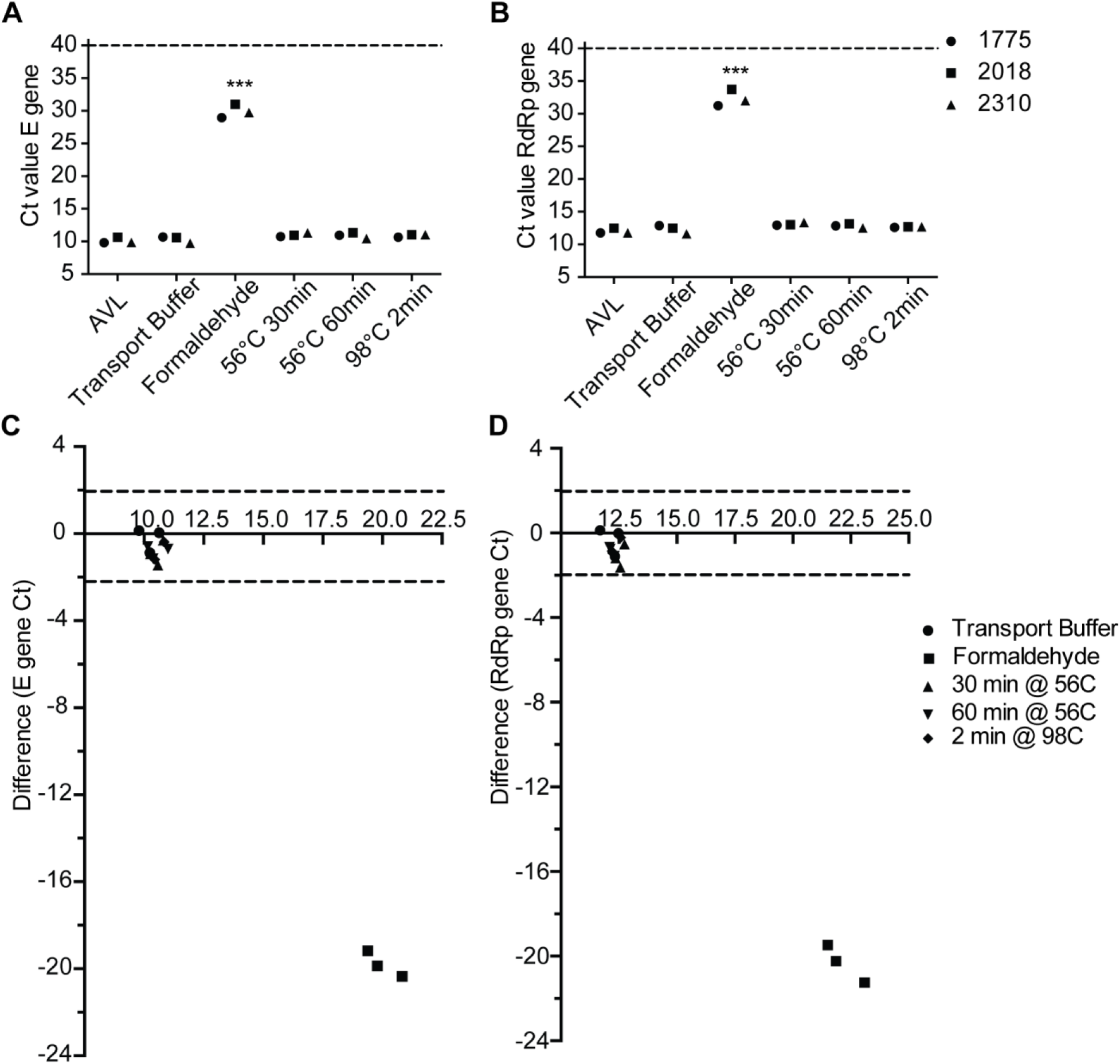
Comparison of Ct values of SARS-CoV-2 (A) E gene, and (B) RdRp gene for three isolates (1775; circles, 2018; squares, 2310; triangles) inactivated by different methods. Inactivation with 2% formaldehyde for 15 min at RT results in significantly elevated Ct values for both genes (***p=0.0001, one-way ANOVA comparison to AVL inactivation). Bland Altman Plots comparing Ct values for (C) E gene, and (D) RdRp gene following inactivation by sample transport buffer (circles), formaldehyde (squares), 30 min at 56°C (triangle), 60 min at 56°C (inverted triangle) and 2 min at 98°C (diamond) compared to AVL. Ct value difference between the two values is plotted on the Y-axis. Cut-off values of 2 and −2 Ct are plotted as dashed lines.

## Discussion

Following the rapid global spread of SARS-CoV-2 and the need for universal testing, more and more individuals are exposed to live virus samples, thereby increasing the chances of occupational infection. The WHO and United States Centers for Disease Control (USCDC) have released laboratory guidelines to mitigate risk of exposure during diagnostic and research procedures [3, 4]. However, despite initial recommendations for handling within contained biosafety cabinets, individuals working with these samples are still required to handle potentially live virus at the initial steps of acquiring and preparing the suspected samples prior testing, thereby increasing the risk of exposure. Potential exposure greatly increases in situations requiring large numbers of samples to be processed under harsh conditions, in underequipped or poorly maintained laboratories, and even within the sample transportation system, such as found in developing or rural areas of the world. Therefore, the continued need for COVID-19 testing worldwide requires utilization of simple and effective inactivation techniques.

Previous studies have been conducted on the effectiveness of chemical inactivation techniques on SARS-CoV-2 [11, 12], the majority of these based on infectious agents of concern such as Ebola [13] and SARS and MERS coronaviruses [14]. As with other viruses, the primary step in the molecular detection of SARS-CoV-2 is viral lysis to begin the extraction of nucleic acids. The buffers used in this lysis step yield varying results [11, 13, 15, 16]; however, unlike previous studies [11], this study found that AVL buffer alone was successfully able to fully inactivate up to 5 log_10_ of virus from three different primary isolates of SARS-CoV-2. Apart from differences in isolates utilized and a slight reduction in titer, it is unclear as to the reasons why AVL buffer fully inactivated in this study versus others, but further work is warranted to determine the exact effectiveness of this step alone.

Inactivating sample transport media, either made in-house or commercially available, also presents an attractive way to inactivate samples at the point of sampling to ensure safe handling along the transport chain and within the laboratory. These inactivating transport media include the key components of many viral lysis buffers including chaotropic agents (GITC), detergents (Triton X-100) and buffering agents (EDTA, Tris-HCL) to inactivate a preserve viral RNA. Previous studies have shown that GITC-lysis buffers are able to inactivate SARS-CoV-2 samples [11, 12]; however, the addition of Triton-X may be necessary for complete inactivation [11]. In line with these studies, commercial sample transport media containing both GITC and Triton-X was successfully able to inactivate up to 5 log_10_ of virus with no loss of molecular diagnostic sensitivity.

Apart from sample media and buffers utilized for diagnostic testing, various disinfectant and inactivating chemicals are available for sample treatment. Formaldehyde has a long history of use for inactivation against a number of viruses and in a number of fixation techniques, including vaccine preparations [17, 18]. Formaldehyde has been shown to successfully inactivate both SARS and MERS [14, 19, 20] and has been suggested to be a viable alternative for disinfection and inactivation of SARS-CoV-2 [19, 20]. Formaldehyde treatment did successfully inactive up to 5 log_10_ of virus; however, this treatment severely impacted viral detection in subsequent molecular testing. This decreased detection is not unexpected as formaldehyde treatment results in RNA degradation and modification [21]. Therefore, formaldehyde treatment does not appear to be a solution for increased molecular SARS-CoV-2 testing; however, it does remain a viable alternative for sample inactivation or disinfection.

Perhaps the most studied technique thus far regarding SARS-CoV-2 has been thermal inactivation at various times and temperatures [11, 22–24]. Several previous studies have shown heat to be an effective inactivation technique against other coronaviruses, including SARS, MERS, and human seasonal strains [14, 23, 25]. Similar to previous studies, 56°C heat treatment for 30 or 60 minutes was fully able to inactivate up to 5 log_10_ of SARS-CoV-2 from three different isolates [11, 22]. Interestingly, while other studies utilized 95°C for 5 to 10 minutes for inactivation, heat treatment at 98°C for only 2 minutes was also able to completely inactivate up to 5 log_10_ of virus. These results are very promising as high heat treatment is extremely rapid and may be a vital addition to the testing arsenal, as RT-PCR can possibly be performed directly from these samples without the need for nucleic acid extraction [26, 27]. Interestingly, the shortened time period of high heat treatment may mitigate some of the reduction in detection seen in previous studies and make this technique more employable [11].

Overall, the agreement and retained sensitivity amongst RT-PCR results, combined with the fact that all methods resulted in 100% virus inactivation up to a viral load of 5 log_10_, suggests that any of the tested methods, except formaldehyde, are useful to inactivate SARS-CoV-2 samples. Given the WHO recommendation to “test, test, test,” these data can help to optimize sample inactivation for austere or remote areas. Indeed, it may be possible to use basic tools such as a stopwatch and boiling water to achieve 100% virus inactivation without compromising sample integrity, significantly decreasing possible exposure during sample transportation and handling, allowing for dissemination of testing to labs with decreased biosafety and biosecurity capacity, and possibly reducing the global demand for a dwindling supply of PPE.

## Acknowledgements

We would like to acknowledge the patients, Cambodian Ministry of Health and Cambodian Center for Disease Control, Rapid Response Team members, and the doctors, nurses, and staff of the reference hospitals in Cambodia for their response to the COVID-19 pandemic. We thank all of the technicians and staff at Institut Pasteur du Cambodge in the Virology Unit and at NAMRU-2 in the Molecular Unit involved in this work.

